# miR167-ARF8, an auxin-responsive module involved in the formation of root-knot nematode-induced galls in tomato

**DOI:** 10.1101/2022.07.29.501986

**Authors:** Yara Noureddine, Martine da Rocha, Jing An, Clémence Médina, Joffrey Mejias, Karine Mulet, Michael Quentin, Pierre Abad, Mohamed Zouine, Bruno Favery, Stéphanie Jaubert-Possamai

## Abstract

- Root-knot nematodes (RKN) from genus *Meloidogyne* induce the dedifferentiation of root vascular cells into giant multinucleate feeding cells. These feeding cells result from an extensive reprogramming of gene expression in targeted root cells, as shown by transcriptomic analyses of galls or giant cells from various plant species.
- Small non-coding RNAs, and messenger RNAs from tomato (*Solanum lycopersicum*) galls and uninfected roots were sequenced. *De novo* microRNA prediction in the tomato genome identified microRNAs expressed in galls and uninfected roots. Statistical analyses identified 174 miRNA genes differentially expressed in galls at 7 and/or 14 days post infection (dpi).
- Integrative analyses combining small non-coding RNA and transcriptome datasets with the specific sequencing of cleaved transcripts identified miRNA targets in tomato galls. Functional analyses of promoter-GUS fusions and CRISPR-Cas9 mutants highlighted the role of the miR167-regulated transcription factor AUXIN RESPONSE FACTOR 8 (ARF8) in giant cell formation.

## Introduction

Root-knot nematodes (RKNs) are major crop pests causing massive yield losses, estimated at millions of Euros annually, worldwide (Blok *et al*., 2008; Abad & Williamson, 2010). These microscopic worms of genus *Meloidogyne* have a wide host spectrum encompassing more than 5,000 plant species, and a wide geographic distribution. After infecting the root, these obligatory parasites induce the *de novo* formation of a specialized feeding site that is crucial for nematode survival. The second-stage RKN juveniles (J2) penetrate the roots and migrate within them; they then inject a cocktail of molecules into five to seven root parenchyma cells (Favery et al., 2016). In response to RKN signals, targeted root parenchyma cells dedifferentiate into giant multinucleate hypermetabolic feeding cells. These “giant cells” form the feeding site supplying the nematode with the nutrients it requires for its development (Favery *et al*., 2020). Dedifferentiation into giant cells involves an initial phase of successive mitoses without cytokinesis, followed a second phase of endoreduplication (de Almeida Engler & Gheysen, 2013). During feeding cell formation, the neighboring cells begin to divide. This whole process results in a swelling of the root to form a gall, the characteristic symptom of RKN infection. Feeding site formation involves several biological processes, including the cell cycle (de Almeida-Engler et al., 2011), metabolic reprogramming (Marella *et al*., 2013), cytoskeleton organization (Caillaud *et al*., 2008), and auxin signaling (Gheysen & Mitchum, 2019). Auxin (indole-3 acetic acid, IAA), is a major plant hormone that plays a key role in root development by regulating cell division and the establishment/maintenance of root primordia (De Smet *et al*., 2007; Weijers & Wagner, 2016). The formation of RKN-induced feeding sites has been shown to involve a peak in auxin levels (Karczmarek *et al*., 2004; Absmanner *et al*., 2013) and gall transcriptome analyses have shown that auxin biosynthesis and auxin-responsive genes are upregulated in *A. thaliana* early galls, whereas the genes encoding repressors of auxin response genes are repressed (Barcala *et al*., 2010).

Multiple transcriptome analyses have been performed on RKN-infected roots, galls or specifically on giant feeding cells, from various plant species, including tomato, initially by microarrays, and more recently by RNA sequencing (Bar-Or *et al*., 2005; Portillo *et al*., 2013; Shukla *et al*., 2018). Four time points in feeding site formation have frequently been investigated in transcriptome analyses: the early phase of feeding site formation at 3 days post infection (dpi), 7 dpi, a time point corresponding to multiple mitoses without cytokinesis, 14 dpi, corresponding to the endoreduplication phase and cell expansion, and, finally, 21 dpi when the feeding cells are mature and fully functional. All these analyses revealed a massive reprogramming of plant gene expression in response to nematode infection, with about 10% of protein-coding genes displaying changes in expression levels in response to RKN infection (Cabrera et al., 2014). However, it remains unclear how this reprogramming occurs and how these genes are regulated.

Small non-coding microRNAs may act as the master regulators of this reprogramming of gene expression (Jaubert-Possamai *et al*., 2019). MicroRNAs are major repressors of gene expression in eukaryotes. Within the plant genome, microRNAs are encoded by *MIR* genes, often organized into multigene families and transcribed as a single-stranded RNA precursor that folds into a typical hairpin structure. This hairpin precursor is processed to generate a microRNA duplex consisting of two complementary strands of 20-22 nucleotides. One of the two strands is then loaded into the ARGONAUTE-1 protein and guides the RNA silencing complex (RISC) to target messenger RNAs through miRNA/mRNA sequence complementarity. The targeting of an mRNA by a microRNA induces its degradation or the inhibition of its translation, depending on the mRNA/miRNA sequence complementarity. Several recent studies have identified microRNAs expressed in galls (Jaubert-Possamai *et al*., 2019) induced by RKN in *Arabidopsis thaliana* (Cabrera *et al*., 2016; Medina *et al*., 2017), in tomato (*Solanum lycopersicum;* Zhao et al., 2015; Kaur et al., 2017), in cotton (*Gossypium hirsutum;* Pan *et al*., 2019; Cai *et al*., 2021) or in rice (*Oryza sativa;* Verstraeten *et al*., 2021). However, the roles of only four microRNAs have been validated by functional analyses: the miR390/tasiRNA/*ARF3* module (Cabrera *et al*., 2016), the miR159/*MYB33* pair (Medina et al., 2017), the miR172/*TOE1/FT* module (Díaz-Manzano *et al*., 2018) in *Arabidopsis* and the miR319/*TCP4* pair in tomato (Zhao et al., 2015).

We investigated the gene regulation network involved in the plant response to RKN through a combination of transcriptome, microRNome and degradome sequencing in uninfected roots of tomato and tomato galls induced by the RKN *M. incognita* at two key time points in gall development: 7 and 14 dpi. We identified 12 miRNA/targeted transcript pairs as robust candidates for the regulation of gall formation. A key role in tomato gall formation was demonstrated for the auxin-responsive miR167/*ARF8* transcript pair in functional analyses.

## Materials and methods

### Biological materials, growth conditions and nematode infection

For *in vitro* experiments, seeds of *Solanum lycopersicum* cv St Pierre or Micro-Tom (wild-type, WT or transgenic *pARF8A::GUS* and *pARF8B::G*US lines (Bouzroud *et al*., 2018)) were surface-sterilized with chlorine solution (44% active chlorine) and washed three times with water. Ten to 15 sterile seeds were sown on a Gamborg B5 (Duchefa Biochemie) agar plates (1x Gamborg B5; pH = 6.4; 1% Sucrose; 0,7% Agar), placed at 24°C for 48 hours for germination, and finally transferred in a growth chamber (8h light; 16h dark, 20°C). *M. incognita* (Morelos strain) J2s were sterilized with HgCl2 (0.01%) and streptomycin (0.7%) as described before (Caillaud & Favery, 2016). One to two weeks after germination, roots were inoculated with 1,000 sterile J2s per petri dishes.

For *in soil* infection assay, *S. lycopersicum* (cv Micro-Tom) WT plants and CRISPR lines (*arf8a, arf8b* and *arf8ab*) were sown and individually transferred in pots filled with a mixture of sand and soil (1:1, vol:vol), kept at 4°C for 48 hours, then transferred in a growth chamber (16h light and 8h dark, at 24°C). Two weeks after germination, each plant was inoculated with 200 J2s. Infection rate was evaluated six weeks after inoculation. The root system of each plant was collected, rinsed with tap water, weighted and stained for 30 s. in eosin (Sigma) solution (0.5%). Galls and egg masses were counted for each root under the binocular magnifier MZFLIII (Leica). Mann–Whitney *U*J tests (α = 2.5%) were performed to determine the significance of the differences in the numbers of egg masses and galls per root observed between mutants and WT.

### BABB clearing

For giant cell area measurements, galls were collected 21 days post-infection (dpi), cleared in benzyl alcohol/benzyl benzoate (BABB) as previously described (Cabrera *et al*., 2018; Mejias *et al*., 2021) and examined under an inverted confocal microscope (model LSM 880; Zeiss). The mean areas of giant cells in each gall, for WT and CRISPR lines, for two biological replicates, were measured with Zeiss ZEN software. The impact of the mutation on the giant cell surface was analysed using a Mann & Whitney Test.

### RNA extraction

Total RNAs, including small RNAs (< 200 nt), were isolated from *in vitro* tomato (St Pierre) galls or uninfected root fragments at 7 and 14 dpi. Approximately 40 galls or uninfected roots devoid meristems were independently frozen into powder by using a tissue lyser (Retsch; MM301) at 30 Hertz frequency for 30 seconds with 4 mm tungsten balls (Retsch; MM301). Total RNAs were extracted from these samples with the miRNeasy Mini Kit (Qiagen), according to the manufacturer’s instructions, with three additional washes in RPE buffer.

### RNA sequencing

Small RNA libraries were generated by ligation, reverse transcription and amplification (11 cycles) from total RNAs (1 µg), with the reagents of the NEB Next Small RNA Library Prep Set for Illumina. Libraries were then quantified with the Bioanalyzer High Sensitivity DNA Kit (Agilent) and sequenced at the Nice-Sophia Antipolis functional genomics platform (France Géenomique, IPMC, Sophia Antipolis, France). The full raw sequencing data were submitted to the GEO database (http://www.ncbi.nlm.nih.gov/geo/), accession number PRJNA799360.

PolyA-RNA libraries from St Pierre tomato were generated from 500 ng of total RNA using Truseq Stranded mRNA kit (Illumina). Libraries were sequenced on a NextSeq 500 platform (Illumina) with 2□×□75bp paired-end chemistry as described in (Mejias *et al*., 2022). RNA-seq data are available at SRA database accession number #PRJNA799360. PolyA sequencing of galls from the *arf8* mutants and microtom wild type was performed by the Beiging Genomics Institute (BGI) by using the DNBSeq technology. RNA-seq data are available at SRA database accession number #XXX

### miRNAome and transcriptome analysis

For each small RNA library, adapters were trimmed and reads matching ribosomal RNA, mitochondrial RNA and repeat sequences were removed by performing Blast analyses with the sequences listed in the Rfam database (Nawrocki et al., 2015). The STAR 2.5 aligner (: --twopassMode Basic --alignEndsType EndToEnd) was then used to align the trimmed reads (Dobin *et al*., 2013) on a virtual concatenated genome generated from the *S. lycopersicum* genome (V3.01, annotation V3.2) and the *M. incognita* genome (Blanc-Mathieu *et al*., 2017). Each read was attributed to the *S. lycopersicum* and/or *M. incognita* genome on the basis of the best alignment obtained. Low-quality mapped reads were removed. The htseq-count package version 0.9.1 (Anders *et al*., 2015) was used to count reads mapping perfectly onto the *S. lycopersicum* genome. The counts for protein coding genes from each replicate were used for differential expression analysis with the R package EdgeR version 3.4.1 (Robinson *et al*., 2009) and DSeq2 (Anders & Huber, 2010). Differentially expressed miRNAs, identified with a false discovery rate of 5% (adjusted pvalue<0.05; Benjamini-Hochberg adjustment).

*De novo* microRNA encoding genes were predicted in tomato genome V3.0 by using three algorithms MirCat (Paicu *et al*., 2017), Shortstack (Axtell, 2013) and MirDeep plant (Yang & Li, 2011) with default parameters. The sequence homology between newly predicted miRNA mature sequences and mature miRNA sequences listed in miRBase 22.1 was analysed by using SSearch algorithm (Kozomara *et al*., 2019). The HTSEQCOUNT package (Anders *et al*., 2015) was used to count reads mapping perfectly onto the predicted *S. lycopersicum* mature microRNA 5P or 3P sequence. Reads mapping to multiple loci were counted for each of the loci concerned. The counts for mature miRNAs (5P and 3P) from each replicate were used for differential expression analysis by using DSeq2 statistical analysis (Anders and Huber, 2010). Mature miRNAs with an adjusted p value below 0.05 were considered as differentially expressed.

GO analyses of genes differentially expressed in galls were performed by using over-representation test from PANTHER analysis tools (Mi *et al*. 2019) with a Fisher’s exact test, a FDR threshold of 0.05 and by selecting “Biological Process » as GO category.

### Degradome analysis

Degradome libraries were constructed from total RNAs extracted from galls at 7 and 14 dpi by Vertis Biotechnologie (Freising, Germany) using the parallel analysis of RNA ends (PARE) protocol described by German *et al*. (2009). The PARE libraries were sequenced on an Illumina High Sequencing 2000 platform. The full raw sequencing data were submitted to the GEO database (http://www.ncbi.nlm.nih.gov/geo/), accession XXX. To identify miRNA targets, degradome reads were analysed and classified by using the CleaveLand 4.0 (Addo-Quaye *et al*., 2009) algorithm with default parameters. All hits are classified into five categories based on the abundance of the diagnostic cleavage tag relative to the overall profile of degradome tags matching the targets.

### GUS staining analysis

We localized the promoter activity in tomatoes transgenic lines expressing a reporter gene GUS fused to the promoter of the two tomato genes *ARF8A* (*Solyc02g037530*) and *ARF8B* (*Solyc03g031970*) (pARF8A:GUS and pARF8B:GUS lines) (Bouzroud *et al*., 2018). We inoculated 21-day-old seedlings *in vitro*, as described above. GUS staining was performed 7 and 14 dpi as previously described (Noureddine *et al*. 2022), and the roots were observed under a Zeiss Axioplan 2 microscope. Stained galls were dissected, fixed by incubation in 1% glutaraldehyde and 4% formaldehyde in 50 mM sodium phosphate buffer pH 7.2, dehydrated, and embedded in Technovit 7100 (Heraeus Kulzer, Wehrheim, Germany), according to the manufacturer’s instructions. Sections were cut and mounted in DPX (VWR International Ltd, Poole, UK), and observed under a Zeiss Axioplan 2 microscope (Zeiss, Jena, Germany).

### Quantitative RT-PCR

Total RNA was extracted from galls and uninfected roots produced in soil with the miRNeasy kit (QIAGEN) according to the manufacturer’s instruction. 500ng of total RNA were subjected to reverse transcription with the Superscript IV reverse transcriptase (Invitrogen). qPCR analyses were performed as described by Nguyen *et al*. (2018). We performed qPCR on triplicate samples of each cDNA from three independent biological replicates. *SlPSKR1* (*Solyc01g008140*) and a gene coding for a Sucrose Synthase (*SuSy3, Solyc07g042550*) were used for the normalization of qRT-PCR data. Quantifications and statistical analyses were performed with SATqPCR (Rancurel *et al*., 2019), and the results are expressed as normalized relative quantities. Primers used to amplify the premature miRNAs and the transcripts are listed in Table S1.

### Generation of *ARF8* mutants by CRISPR/Cas9

For CRISPR/Cas9 construct, the sgRNA sequence (AAGCTTTCAACATCAGGAA) commune to SlARF8a and SlARF8b was designed by using the CRISPR-P website tool (http://cbi.hzau.edu.cn/crispr/). The sgRNA was cloned into pAGM4723 final vector by golden gate ligation method. Construct was confirmed by sequencing before introduction into the C58 *Agrobacterium tumefaciens* strain. Tomato seedlings were used for the next step plant transformation according to Hao et al. (2015).

## Results

### Gall formation results from a massive reprogramming of gene expression in root cells

Two statistical methods, DSeq2 and EdgeR, were used to compare transcript levels between tomato galls and uninfected roots. These two methods were applied to 19,918 genes, and those found to be differentially expressed by both methods, with an adjusted *p*-value below 0.05, were identified as differentially expressed genes (DEG). We found 1,958 DEGs at 7 dpi **(Table S2)** and 3,468 DEGs at 14 dpi **(Table S3 and Figure 1a)**. In total, 1,239 genes were identified as DEGs at both 7 and 14 dpi, including 625 genes downregulated and 600 genes upregulated at both time points and 14 DEGs with opposite patterns of change in expression levels at 7 and 14 dpi. The 719 genes displaying differential expression in galls specifically at 7 dpi comprised 327 upregulated and 392 downregulated genes. The 2,229 genes displaying differential expression in galls specifically at 14 dpi comprised 1,006 upregulated and 1,223 downregulated genes. The change in gene expression in galls detected by sequencing was confirmed by RT-qPCR for six of these genes at 7 dpi and six at 14 dpi (**Figure S1 and Table S4**). Gene ontology (GO) analysis DEGs in galls at 7 and/or 14 dpi revealed an overrepresentation of genes associated with biological processes previously reported to be involved in the formation of giant cells **(Table S5)**, including i) “cell division”, with multiple categories linked to cytokinesis and cell wall biogenesis, ii) “response to auxin”, iii) “response to endogenous stimulus” (including response to hormone and to cytokinin) and iv) “response to abiotic stress”. This analysis confirms that a massive reprogramming of gene expression in root cells underlies the formation of galls and feeding cells.

**Figure 1.**
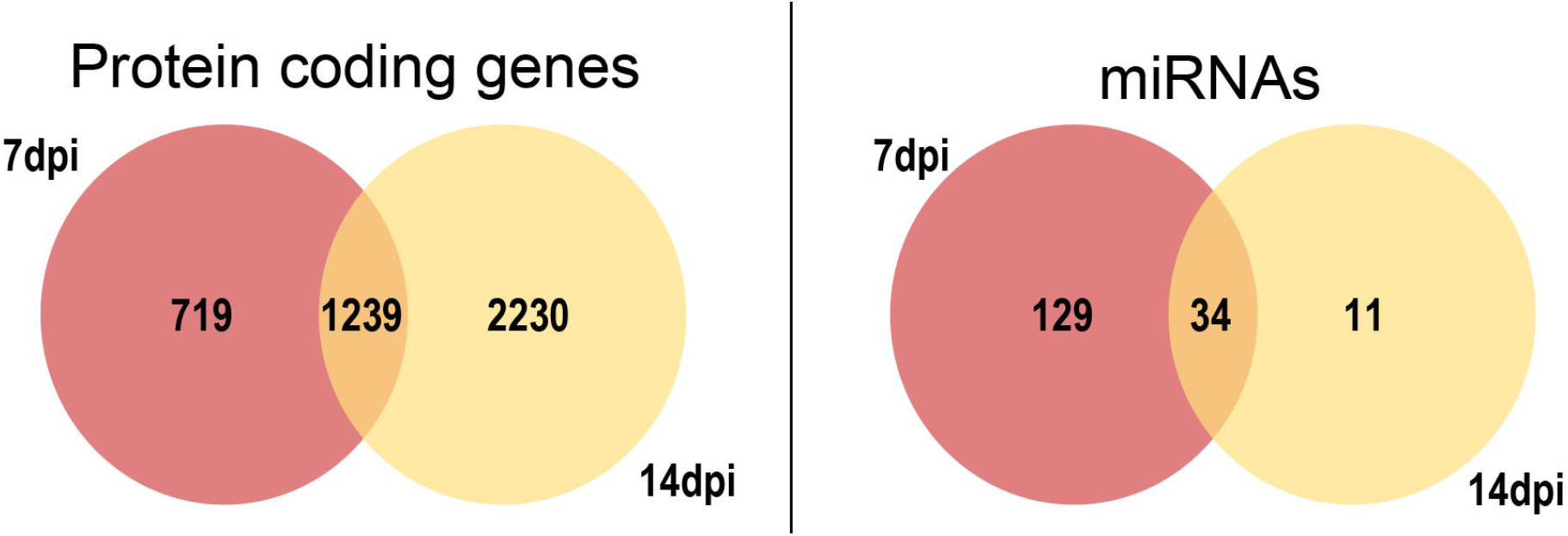
Tomato protein-coding genes and miRNAs differentially expressed in *M. incognita*-induced galls relative to the corresponding uninfected roots, at 7 and/or 14 dpi. The numbers of genes differentially expressed at 7 days post inoculation (dpi) (pink) and/or 14 dpi (yellow) between galls and the corresponding uninfected roots are indicated in Venn diagrams.

### microRNAs regulate gene expression in galls

For the identification of regulators of gene expression in galls, we constructed libraries of small non-coding RNAs from tomato galls and uninfected roots, from three independent replicates at two points of gall development: 7 and 14 dpi. These libraries were sequenced, generating a total of 333,949,327 raw reads **(Table S6)**. The reads were cleaned and mapped to a virtual genome constructed from the *S. lycopersicum* genome (genome V3.0; ITAG3.3) concatenated with the genome of *M. incognita* (genome V2.0; Blanc-Mathieu et al., 2017) to reflect the dual composition of root galls. A *de novo* prediction of microRNAs was then performed (**Table S7**), based on an integration of the results of three prediction algorithms: MirCat, Shortstack and MirDeep plant.

Levels of microRNA expression were compared between galls and uninfected roots in DSeq2 statistical analyses (Anders & Huber, 2010). We identified 174 mature microRNAs (5P and/or 3P) corresponding to 148 *MIR* genes as differentially expressed (DE) between uninfected roots and galls at 7 and/or 14 dpi **(Table S8)**. We identified 129 of the 174 mature microRNAs DE in galls as specifically DE at 7 dpi, 11 as specifically DE at 14 dpi and 34 mature microRNAs were found to be DE in galls at both 7 and 14 dpi **(Figure 1b)**. These 148 *MIR* genes DE in galls comprised 65 known *MIR* genes listed in miRbase (Kozomara *et al*., 2019) and 73 previously unknown *MIR* genes. The 65 known *MIR* genes DE in galls are organized into 20 miRNA families, 14 of which are conserved between tomato and other plants, the other six being specific to tomato.

### Integration of data from the transcriptome, small RNAs and degradome sequencing to construct a gene-microRNA regulation network in galls

Once the miRNAs expressed in galls had been identified, the transcripts cleaved by the microRNAs in galls were identified by degradome sequencing (German *et al*., 2009) on mRNA extracted from galls at 7 (G7) and 14 dpi (G14). The CleaveLand pipeline (Addo-Quaye *et al*., 2009) was used to analyze degradome sequencing data and to predict the mRNAs cleaved by miRNAs in galls. We restricted our analysis to the highest confidence targets by selecting CleaveLand categories 0 and 1 with a degradome *p*-value below 0.05. In total, 153 transcripts targeted by microRNAs in galls were identified **(Table S9)**, including 58 targets common to both the G7 and G14 libraries, whereas 45 targets were identified specifically in the G7 library and 50 were found only in the G14 library. We identified 111 targets of 135 known miRNAs from 39 known miRNA families. The 298 newly identified miRNAs in galls included 46 that targeted 47 transcripts in galls at 7 and/or 14 dpi.

We integrated transcriptome, microRNA and degradome sequencing data to construct a a gene-miRNA regulation network putatively involved in the gall formation. Transcriptome analysis showed that 32 of the 153 transcripts identified as targeted by microRNAs expressed in galls were DE in galls. Nineteen of the targeted genes were DE in galls at both 7 and 14 dpi; 11 of these genes were upregulated and eight were downregulated. Five targeted genes were specifically DE at 7 dpi, including three transcripts that were upregulated and two that were downregulated in galls. At 14 dpi, only eight transcripts identified as targets were DE, three of which were upregulated, the other five being downregulated. Most plant miRNAs silence gene expression by cleaving the targeted transcripts. An inverse correlation of expression profiles between the microRNA and its target gene is, therefore, usually expected. We identified 12 miRNA/mRNA pairs for which such an inverse correlation of expression levels was observed **(Table 1)**. These miRNA/mRNA pairs are the most robust candidates for involvement in gall formation.

**Table 1.**
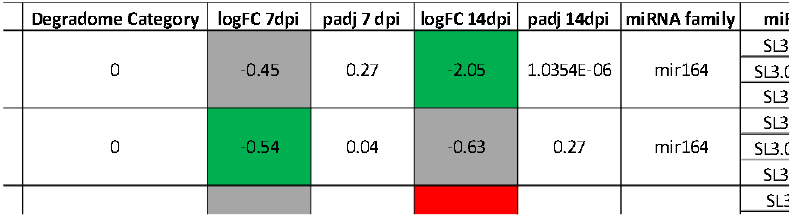
The 12 miRNA/mRNA pairs with inversely regulated expression profiles. Degradome analysis identified 12 genes targeted by miRNAs in galls. *Solanum lycopersicum* (Solyc) gene expression levels were compared between galls and uninfected roots, by two statistical methods (DSeq2 and EdgeR), and the expression of mature miRNAs was compared by DSeq2. Gall/root fold-change differences in expression (LogFC) at 7 and 14 days post infection (dpi) and the adjusted *p*-value obtained by the Benjamini-Hochberg method (adj *p*-value) are indicated for genes and miRNAs. The genes and mature microRNAs upregulated in galls are shown in red, and those downregulated in galls are shown in green.

### ARF8 auxin-related transcription factors are expressed in nematode-induced feeding sites

Among the 12 microRNA/mRNA pairs with opposite patterns of gene expression, two of the strongest genes candidates for regulation by a microRNA are the *AUXIN RESPONSE FACTORS 8A* (*Solyc02g037530*) and *8B* (*Solyc03g031970*), both of which are cleaved miR167. These two genes are ARF transcription factors, which relay auxin signaling at the transcriptional level by regulating the expression of auxin-responsive genes (Guilfoyle & Hagen, 2007). Transcriptomic analyses of galls showed that *ARF8B* was overexpressed in tomato galls at 7 and 14 dpi and *ARF8A* was overexpressed at 14 dpi **(Table 1)**.

Five *MIR167* genes were identified in the tomato genome, including the four described by Liu *et al*. (2014), all of which have the same mature sequence and are downregulated in galls, whereas both the *ARF8* genes were found to be upregulated. The fifth *MIR167* gene was annotated as SLYMIR167B in miRBase (Kozomara *et al*., 2019) and was not DE in galls. However, it was expressed at a much lower level than the other four *MIR167* genes (**Figure S2**). *ARF8B* transcripts have been shown to be cleaved by miR167 in tomato (Liu *et al*., 2014) and this regulation in conserved in *A. thaliana* (Wu *et al*., 2006). The downregulation of miR167 and the upregulation of *ARF8A* and *ARF8B* observed in galls suggest that, by repressing *MIR167* expression, the RKN prevents the *ARF8* silencing by miR167 that occurs in uninfected roots.

We investigated the spatiotemporal expression of *ARF8A* and *ARF8B* in RKN-infected roots *in vivo* further, by analyzing the activity of both the *ARF8A* and *ARF8B* promoters in transgenic tomato lines expressing promoter-GUS fusions (Bouzroud *et al*., 2018). Strong blue staining indicating GUS activity was observed 7 and 14 dpi in galls from two independent *pARF8B::GUS* and *pARF8A::GUS* lines and in root tips from uninfected roots **(Figure 2a-f)**. Histological sections of the galls showed strong GUS staining within the giant feeding cells and in neighboring cells (NC) at 7 and 14 dpi for both *ARF8A* and *ARF8B* lines **(Figure 3a-d)**. The strong activity of both the *ARF8A* and *ARF8B* promoters observed in galls *in vivo* confirms the upregulation in galls observed on transcriptomic analysis.

**Figure 2.**
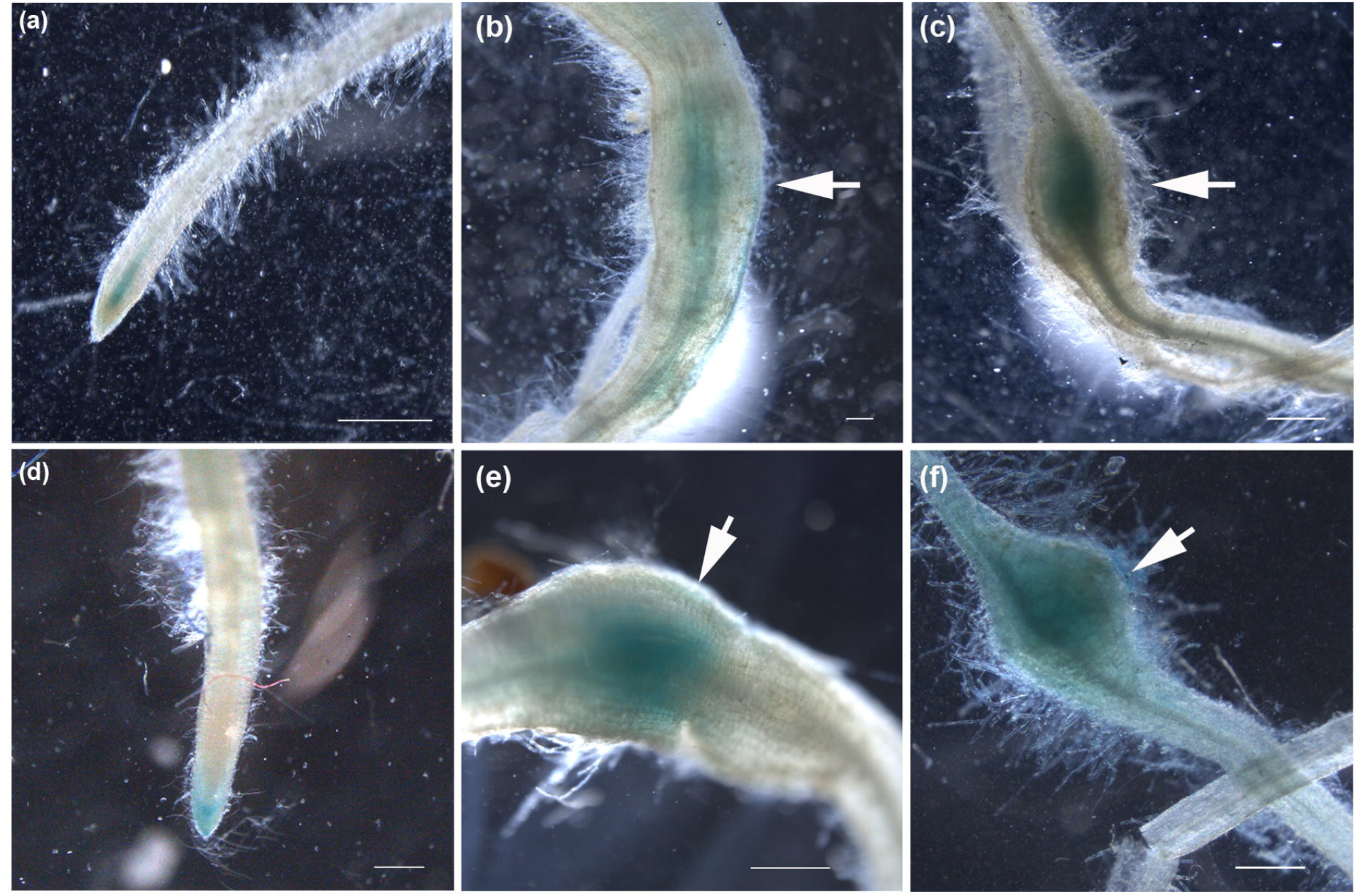
*ARF8A* and *ARF8B* are strongly transcribed in tomato galls induced by *M. incognita*. The activity of the *ARF8A* and *ARF8B* promoters (*pARF8A* and *pARF8B*) was studied in galls induced by *M. incognita* in *S. lycopersicum* lines expressing the *pARF8A::GUS* or the *pARF8B::GUS* construct, at 7 and 14 days post inoculation (dpi). Blue staining indicating GUS activity under the control of *pARF8A* was observed in (a) uninfected root tips and (b-c) galls induced by *M. incognita* at 7 dpi (b) and 14 dpi (c). GUS activity under the control of *pARF8B* was observed in (d) uninfected root tips and in (e-f) galls at 7 dpi (e) and 14 dpi (f). Bars: 500 µm.

**Figure 3.**
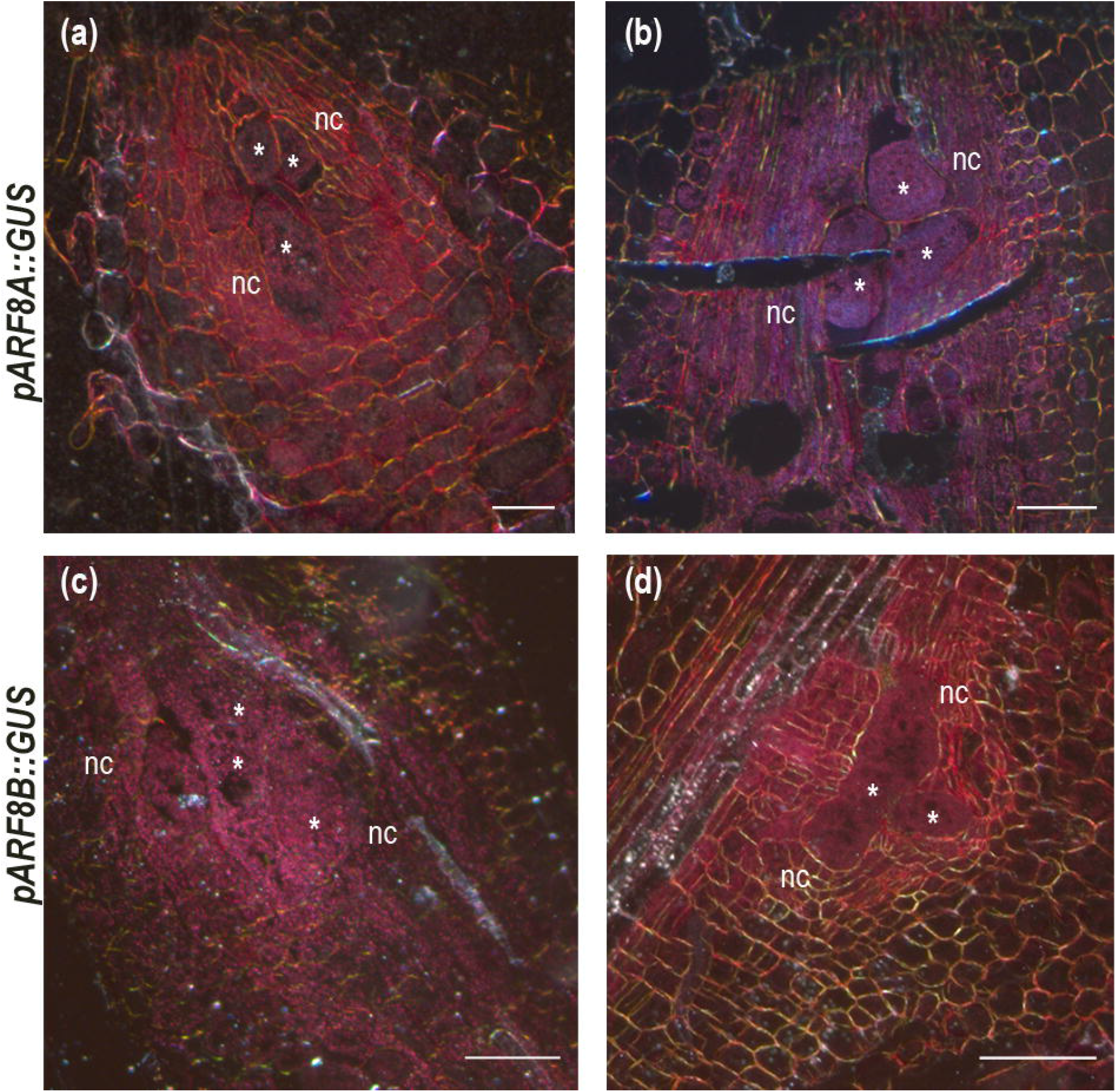
*ARF8A* and *ARF8B* are strongly transcribed in 14 dpi nematode-induced feeding sites. The activity of the *ARF8A* and *ARF8B* promoters (*pARF8A* and *pARF8B*) was studied in galls induced by *M. incognita* in *S. lycopersicum* expressing the *pARF8A::GUS* (a-b) or the *pARF8B::GUS* (c-d) construct, at 7 and 14 days post inoculation (dpi). Gall sections (5 µm) were cut after GUS staining and observed by dark-field microscopy. Red staining, reflecting GUS activity, was observed in giant cells and neighboring cells in *pARF8A::GUS* galls (a) 7 dpi and (b) 14 dpi. Strong GUS activity was observed in giant cells and neighboring cells in the galls of *pARF8B::GUS* plants at (c) 7 dpi and (d) 14 dpi. *, giant cells; nc: neighboring cells; Bars: (a,b) 100 µm (c,d) 50 µm.

### Generation of *SlARF8A* and *SlARF8B* mutants by the CRISPR/Cas9 gene-editing system

We investigated the function of *SlARF8A* and *SlARF8B* during plant-nematode interaction, by generating tomato Micro-Tom *slarf8* KO mutants with CRISPR/Cas9 gene-editing technology. *SlARF8A* and *SlARF8B* single and double mutants were obtained with a sgRNA complementary to a region identical in both genes (Figure 4a). The transformed lines were screened and six independent R0 lines were generated and validated for the presence of the construct in their genome. All mutants had mutations in the targeted region and the features of the mutant plants were similar. In the R1 and R2 generations, the presence of mutation was confirmed in more progeny lines. Three Cas9-free and homozygous mutant lines containing single (*SlARF8A* or *SlARF8B)* or double mutation (*SlARF8A* and *SlARF8B)* were selected in the following experiments. These three mutants types can be classified as *SlARF8A* single mutation (*slarf8acr*), *SlARF8B* single mutation (*slarf8b-cr*) and *SlARF8A&B* double mutation (*slarf8a&b-cr*) (Figure 4b). As shown in figure 4, the deletion mutations led to a frame shift mutation followed by an early stop codon leading to the expression of truncated SlARF8 proteins that do not contain the ARF family functional domains B3, III and IV. For *slarf8a-cr*, a deletion of 2 nt was detected in the sgRNA1 targeted region, leading to a 13 amino acid (aa) protein rather than the 845 aa WT protein; for *slarf8b-cr*, an 11nt deletion was observed in the sgRNA1 target region, leading to a 9 aa protein sequence rather than 843 aa in WT protein; and, for *slarf8a&b-cr*, there was a 2 nt deletion on *SlARF8A* and a 4 nt deletion on *SlARF8B* resulting in a 13 aa SlARF8A protein and a 20 AA SlARF8B protein.

**Figure 4.**
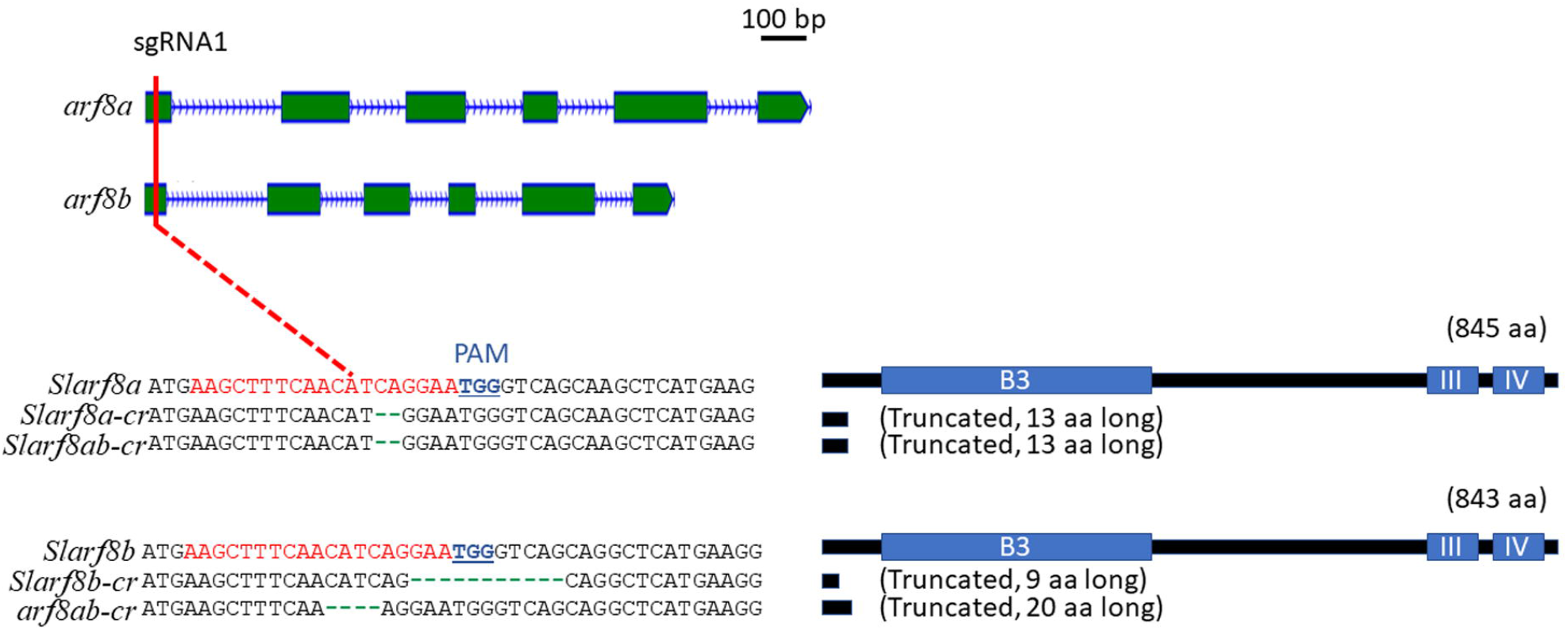
Tomato *slarf8-KO* lines. Generation of slarf8-KO mutant lines by CRISPR/Cas9. Guide RNAs (sgRNA, red bar) anchored next to the Zinc Finger Motif (ZFM) were designed for CRISPR/Cas9 strategy. Protospacer adjacent motif (PAM) are indicated in blue. Mutations within *slarf8* coding sequences corresponding to nucleotide deletions are shown in green. Three types of mutants predicted to produce heavily truncated proteins were chosen for further phenotypic characterization. These mutants are annotated as *arf8a-cr* (*arf8a* single mutant), *arf8a-cr* (*arf8b* single mutant), *arf8ab-cr* (*arf8a* and *arf8b* double mutant). The predicted mutated proteins are schematically illustrated (right panel).

### ARF8 auxin-related transcription factors are involved in tomato-RKN interactions

We investigated the role of both *ARF8A* and *ARF8B* in gall development, by analyzing the effect of CRISPR deletions within *ARF8* coding sequences on *M. incognita* infection. The *arf8a*^*CR-2*^, *arf8b*^*CR-11*^ and *arf8ab*^*CR-2,4*^ double-mutant CRISPR lines had root phenotypes identical to that of the WT **(Figure S3 and Table S10)**. The rate of infection of these CRISPR lines after inoculation with *M. incognita* was determined by counting the galls on infected roots and the egg masses produced by the adult females at the root surface at the end of the RKN lifecycle. A significant large decrease, by approximately 50%, in the numbers of galls and egg masses was observed for the *arf8a*^*CR-2*^, *arf8b*^*CR-11*^ and *arf8ab*^*CR-2,4*^ lines relative to WT plants **(Figure 5a and Table S10)**. Thus, *ARF8* disruption decreases suceptibility to infection, thereby demonstrating that *ARF8A* and *ARF8B* play a key role in the plant-RKN interaction. We investigated the reasons for the lower susceptibility of the *arf8a*^*CR-2*^, *arf8b*^*CR-11*^ and *arf8ab*^*CR-2,4*^ lines, by measuring the area covered by giant cells directly with a confocal microscope, after gall clearing with BABB (Cabrera *et al*. 2017). A comparison of the mean surface areas covered by giant cells in each gall showed that giant cells from the CRISPR lines were approximately 30% smaller than those from control plants **(Figure 5b-c)**. These results demonstrate that the expression of *ARF8A* and *ARF8B* is required for correct giant cell development during the tomato-RKN interaction.

**Figure 5.**
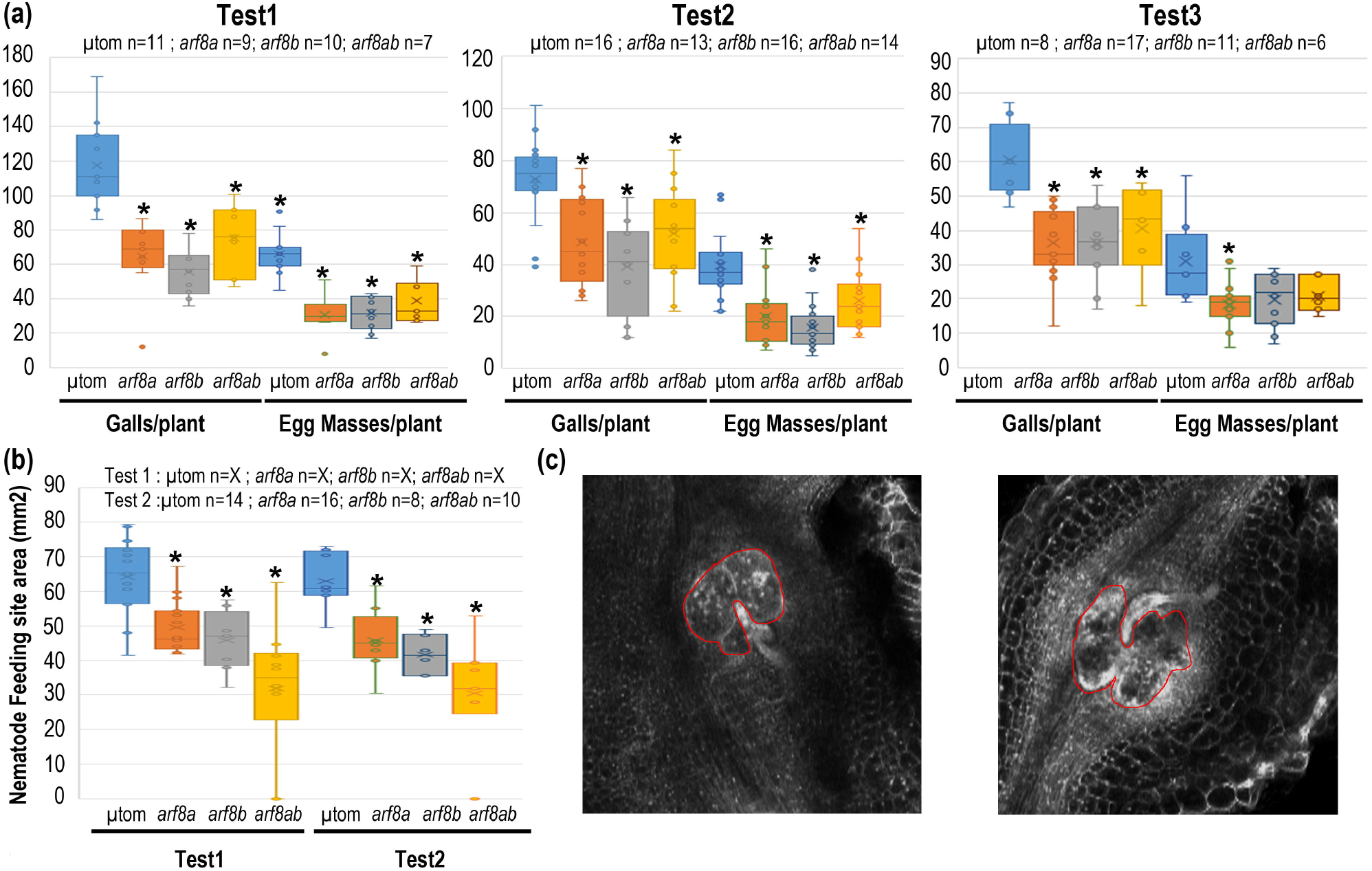
The single mutants *arf8a*^*CR-2*^ and *arf8b*^*CR-11*^ and the double mutant *arf8ab*^*CR-2,4*^ were significantly less susceptible to *M. incognita* than the wild type. (a), The susceptibility of the single and double CRISPR-Cas9 mutant lines and wild-type (WT) MicroTom plants to *M. incognita* was evaluated by counting the numbers of galls and egg masses per plant (G/plant and EM/plant, respectively) in two independent infection assays in soil. (b) The effect of deleting *ARF8A* and/or *ARF8B* on the development of giant feeding cells was further evaluated by measuring the size of the feeding site produced in each mutant line and comparing it to that in the WT. Galls were collected 21 days post infection (dpi) *in vitro* to measure the area (mm^2^) covered by the giant cells by the BABB clearing method (Cabrera et al., 2018). The impact of plant genotype was assessed in Mann-Whitney tests. *, *P* < 0.05. Boxes indicate the interquartile range (25th to 75th percentile). The central lines within the boxes represent the medians. Whiskers indicate the minimum and maximum usual values present in the dataset. The circle outside the box represents an outlier. n, the number of plants analyzed in each assay. Bars 50 µm.

### Identification of *ARF8A-* and *ARF8B-*regulated genes in galls

For the identification of genes regulated by ARF8A and ARF8B in galls, mRNA from 14 dpi galls of *arf8a*^*CR-2*^, *arf8b*^*CR-11*^ mutants and WT Micro-Tom tomatoes were sequenced. Transcript levels in the galls from WT and mutant plants were compared in DESeq2 and EdgeR statistical analyses. These two methods identified 189 and 66 genes, respectively, as differentially expressed between galls from WT and *arf8a*^*CR-2*^ or *arf8ab*^*CR-2,4*^ mutants (**Table S11a and b**). Several auxin-inducible genes have already been identified as candidate genes downstream from ARF8s in tomato floral tissue (Liu *et al*. 2014): *SMALL AUXIN-UPREGULATED RNAs* (SAUR; *Solyc07g042490*.*1*.*1*) and two *EXPANSINS* (*Solyc08g077900*.*3*.*1* and *Solyc08g077910*.*3*.*1*) were found to be repressed in *arf8a*^*CR-2*^tomato galls. Only 16 DEGs were common to both mutants (**Table S11c**). These genes are located directly or indirectly downstream from ARF8 in tomato galls. No defense marker genes, such as orthologs of salicylic acid-mediated response marker *PATHOGENESIS-RELATED PROTEIN-1* (PR1), jasmonic acid-mediated defense maker *PROTEINASE INHIBITOR 2* (*PIN2*) or *PLANT DEFENSINs*, were found to be induced in *arf8a*^*CR-2*^ or *arf8ab*^*CR-2,4*^ galls. This absence of defense marker gene induction in the galls of *arf8a*^*CR-2*^ and *arf8ab*^*CR-2,4*^ mutants indicates that the decrease in galls and egg masses observed is due to a loss of susceptibility rather than the induction of a plant defense mechanism. Together with the requirement of ARF8s for correct giant cell development, this findings supports a key role for ARF8s in feeding site formation.

## Discussion

### Identification of miRNA/mRNA target pairs involved in gall formation

RKN of the genus *Meloidogyne* are highly polyphagous sedentary plant parasites that can induce the formation of giant feeding cells in most crop species. The formation of feeding cells by RKN has been shown to result from a massive reprogramming of plant gene expression induced by the nematode (Jammes *et al*., 2005; Fuller *et al*., 2007; Ibrahim *et al*., 2011; Damiani *et al*., 2012; Yamaguchi *et al*., 2017; Kaur *et al*., 2017). In this study, we identified 4187 protein-coding genes, corresponding to 12.3% of all annotated tomato genes, as differentially expressed in tomato galls 7 and 14 dpi with *M. incognita* relative to uninfected roots. This proportion is consistent with previous transcriptomic analyses in *Arabidopsis* and tomato (Yamaguchi *et al*. 2017; Portillo *et al*. 2013). We investigated the regulators of this reprogramming of gene expression, by analyzing the expression of microRNAs and mRNAs in galls and uninfected tomato roots and using degradome sequencing to identify transcripts cleaved under the guidance of microRNAs. Finally, a gene regulation network for gall development was built by integrating all these -omics data. Twelve of the 153 transcripts identified as targeted by microRNAs in tomato galls in degradome analysis were considered the most robust candidates, based on their expression profiles, which were inversely correlated with those of the corresponding microRNA. Some of these 12 miRNA/mRNA pairs have already been reported to be involved in the plant-nematode interaction in *Arabidopsis:* the miR408/UCCLACYANINE (blue copper protein) and the miR398/copper superoxide dismutase (Noureddine *et al*. 2022), and the auxin-regulated miR172/APETALA2 pair (Díaz-Manzano *et al*. 2018). Moreover, other pairs as also appear to be interesting candidates based on the processes in which they are involved. This is the case, for example for the miR164/NAC transcription factor involved in lateral root development (Guo *et al*. 2005). In *Arabidopsis* root, the auxin-mediated induction of miR164 induces the silencing of *NO APICAL MERISTEM-1 (NAC1)* transcripts, thereby affecting transmission of the auxin signal and regulating lateral root growth.

### *ARF8s* are regulated at posttranscriptionally by miR167 in tomato roots

Among the 12 most robust miRNA/mRNA pairs identified as putatively involved in the formation of galls, the *ARF8A* and *ARF8B* targets of miR167 were selected for further functional analyses based on their role in auxin signaling, as auxins are a class of hormones controlling root development and architecture (De Smet *et al*., 2007; Quint & Gray, 2008; Majda & Robert, 2018). ARF8 is an auxin-responsive factor (ARF). The transcription factors of this family regulate the activation or repression of auxin-induced genes by binding to the auxin response elements (AuxREs) in their promoters (reviewed in Guilfoyle and Hagen, 2007; Chandler, 2016; Li et al., 2016). The ARF gene family is conserved across the plant kingdom and is well described in various plant species, including *A. thaliana* (23 genes) (Hagen & Guilfoyle, 2002), *S. lycopersicum* (22 genes) (Zouine *et al*., 2014), and *Oryza sativa* (25 genes) (Wang *et al*., 2007). The ARF family has been implicated in the regulation of plant developmental processes, such as embryo morphogenesis (Rademacher *et al*., 2011), the formation of lateral roots in response to low levels of nitrogen (Gifford *et al*. 2008), the formation of adventitious roots (Lee *et al*., 2019), leaf structure and senescence (Wilmoth *et al*., 2005), flower development (Ellis *et al*., 2005) and fruit initiation (Liu *et al*., 2014). Like ARF5, 6, 7 and 19, ARF8 has been described as a transcriptional activator (reviewed in Guilfoyle and Hagen, 2007). In *Arabidopsis*, a partial redundancy between ARF8 and ARF6 has been reported, with both these activators silenced by miR167 (Reeves *et al*. 2012). The *Arabidopsis arf6arf8* double mutant has defective flower development, as the flower is entirely sterile (Nagpal, 2005), whereas the *arf8* mutant presents defects of pollination and fertilization (Tian *et al*., 2004; Vernoux *et al*., 2011). A role for ARF8 has been reported in the formation of lateral roots in *Arabidopsis* and soybean (Gifford *et al*., 2008; Wang *et al*. 2015) and in the formation of adventitious roots (Gutierrez *et al*., 2009). ARF6 and ARF8 were recently implicated in cambium establishment and maintenance (Ben-Targem *et al*., 2021). The *arf6arf8* double mutant displays low levels of xylem occupancy and an absence of fiber accumulation until very late stages of plant growth.

In *A. thaliana* and tomato, *ARF8* genes are regulated posttranscriptionally by miR167 (Wu *et al*., 2006; Liu *et al*., 2014). Transcriptomic analyses of galls showed that *ARF8B* is overexpressed in tomato galls at 7 and 14 dpi, whereas *ARF8A* is overexpressed at 14 dpi. The infection of tomato lines expressing the GUS reporter gene under the control of the *ARF8A* or *ARF8B* promoter revealed high levels of activity for both *ARF8* promoters in giant cells and neighboring cells at 7 and 14 dpi, confirming the overexpression observed in the transcriptomic analyses. Gall degradome analysis identified *ARF8A* and *ARF8B* transcripts as cleaved by members of the miR167 family. Four tomato *MIR167* genes encode mature proteins with identical sequences and are downregulated in tomato galls at 7 and 14 dpi. This suggests that the *ARF8A* and *ARF8B* transcripts are cleaved by miR167 in uninfected tomato roots, as demonstrated in *A. thaliana* roots. We showed that RKN infection induces the inhibition of miR167 in galls, thereby decreasing the cleavage of *ARF8* transcripts by miR167, resulting in an overexpression of *ARF8A* and *ARF8B* in galls.

### Auxin is a major factor regulating the formation of feeding cells in tomato

We used tomato lines with CRISPR deletions within the *ARF8A, ARF8B* and *ARF8AB* coding sequences to analyze the function of *ARF8* in plant-nematode interactions. The *arf8a, arf8b* and *arf8ab* lines displayed decreased susceptibility to nematode infection, with fewer gall and egg masses in mutants than in wild-type tomato plants. Moreover, the phenotyping of giant feeding cells within cleared galls showed the giant cells from the three CRISPR lines to be smaller than those from the wild type. These defects, associated with CRISPR mutations, confirmed the involvement of *ARF8A* and *ARF8B* in the tomato response to RKN interaction and that requirement for functional ARF8A and ARF8B for the correct development of feeding cells.

MicroRNAs and transcription factors regulate *ARF8* expression, whereas auxin peaks regulate ARF8 activity: when auxin levels are high, ARF8 (class II ARFs) activates the transcription of auxin-responsive genes (Tiwari *et al*., 2003). Auxin is known to be a major factor regulating the formation of feeding cells in response to RKN signals (reviewed in Gheysen and Mitchum, 2019; Oosterbeek *et al*., 2021). Microarray analyses of *A. thaliana* gall transcripts have revealed an early activation of genes responsible for auxin homeostasis and auxin-responsive genes, and a downregulation of repressors of auxin responses (Hammes *et al*., 2005; Jammes *et al*., 2005; Barcala *et al*., 2010). Studies of lines expressing reporter genes under the control of the synthetic auxin-responsive DR5 promoter showed that this promoter is activated in galls induced by RKN (Hutangura *et al*., 1999; Karczmarek *et al*., 2004; Absmanner *et al*., 2013b). In *A. thaliana* galls, a strong signal was detected in both giant cells and neighboring cells at 4 dpi for DR5:GUS lines (Cabrera *et al*., 2014). This increase in auxin levels in the galls may be controlled by either the plant or the nematode. Auxin-mimicking compounds have been found in nematode secretions (De Meutter *et al*., 2003, 2005).

### ARF8 is involved in plant responses to biotic and abiotic stresses

ARF8 transcription factors have been implicated in plant responses to microorganisms. *ARF8* is regulated in tomato leaves in response to biotic stresses, such as flagellin treatment or infection with *Pseudomonas syringae* (Bouzroud *et al*., 2018). A recent study in *Arabidopsis* provided evidence for a suppression of miR167 expression, together with an induction of *ARF6* and *ARF8*, in response to infection with *P. syringae* pv. tomato DC3000 in *A. thaliana* (Caruana *et al*., 2020). The P35S:*MIR167* and *arf6 arf8* double mutants were found to be more resistant to bacterial infection than the wild type. The authors suggested that ARF6 and ARF8 modulate salicylic acid defenses response to *P. syringae* infection under miR167 regulation. Furthermore, soybean ARF8A and ARF8B have been shown to downregulate the nodulation induced by miR167 (Wang et al., 2015). All these studies suggest that the auxin-responsive pathway, including miR167/ARF8, is a key actor in the response to microorganisms. Analysis of the transcriptomes of two tomato CRISPR *arf8* mutants showed no induction of genes associated with plant defense in galls from *arf8* mutants. This finding supports the notion that the lower levels of RKN infection observed in *arf8* mutants result from a loss of susceptibility rather than an enhancement of plant defense.

### ARF8, an environmental hub connecting development and stress responses

We have shown that *ARF8* genes are posttranscriptionally regulated by miR167 in galls, but the transcriptional regulation of these genes has yet to be deciphered. *ARF8* was recently shown to be regulated by a complex network of multiple activating and repressing transcription factors in *A. thaliana* (Truskina *et al*., 2021). Interestingly, some of these transcription factors, such as WUSCHEL, Squamosa Promoter Binding Protein Like-13 (SPL13) and WRKY33, have also been implicated in plant development, and many *ARF8* regulators are also associated with plant responses to biotic and abiotic stresses. Based on these results, Truskina *et al*. suggested that ARF8 may act as an environmental hub connecting development and stress responses mediating auxin responsiveness. The formation of RKN-induced feeding sites interferes with plant developmental processes, including, in particular, the development of lateral roots (Cabrera *et al*., 2014, Olmo *et al*. 2020), which suggests that the nematode may hijack this process. For example, the transcription factor LBD16 and the microRNA miR390a, two key components of the auxin pathway and transducers of lateral root development, are involved in gall formation in *Arabidopsis* and tomato (Cabrera *et al*., 2014, Olmo *et al*. 2020). *ARF8* has also been implicated in lateral root formation in *Arabidopsis* (Gifford *et al*. 2008) and may, therefore, integrate biotic stress and developmental processes during the formation of giant feeding cells. The common induction, in tomato and *Arabidopsis* galls, of *ARF8* and of the transcription factor *LBD16* (Olmo *et al*. 2020) and miR390a (Diaz Manzano *et al*. 2016), suggests that there may be a conserved auxin-mediated molecular pathway in galls. Early in the development of galls in *Arabidopsis*, at 3 dpi, the silencing of *ARF3* by miR390 via tasiRNAs, and the induction of *ARF5* have been shown to be required for parasitism (Cabrera *et al*. 2014a; Olmo *et al*. 2020). These results suggest that there is a complex network involving ARFs and microRNAs responsible for mediating auxin signaling during the development of galls induced by RKN.

## Acknowledgments

The microscopy work was performed at the SPIBOC imaging facility of Institut Sophia Agrobiotech. We thank Dr Olivier Pierre and the entire team of the platform for assistance with microscopy. This work was funded by the INRAE SPE department and the French Government (National Research Agency, ANR) through the ‘Investments for the Future’ LabEx SIGNALIFE: program reference #ANR-11-LABX-0028-01 and IDEX UCAJedi ANR-15-IDEX-0, and by the French-Japanese bilateral collaboration programme PHC SAKURA 2019 #43006VJ. Y.N. was supported by a doctoral fellowship from Lebanon (Municipal Council of Aazzée).

## Author contributions

Y.N., B.F and S.J.P. designed the study, performed the experimental work and wrote the manuscript. Y.N. and C.M. produced biological material for sequencing. M.dR. analyzed NGS data. M.Z. and JA designed, generated and characterized *arf8* CRISPR-Cas9 mutants. KM participated to the qPCR analyses. M.Q., J.M. and P.A. participated in the writing of the manuscript. All the authors analyzed and discussed the data.

## Data avaibility

s

## Supporting Information

**Fig. S1. Quantification of tomato transcripts by RT-qPCR in galls relative to uninfected tomato roots, at 7 and 14 days post infection**

**Fig. S2. Expression of the Sly-miR167 family in galls and uninfected tomato roots.**

**Fig S3. Root phenotype of *crispr arf8* mutants**

**Table S1. Primers used for RT-qPCR**.

**Table S2. Protein-coding genes DE in galls at 7 dpi**

**Table S3. Protein-coding genes DE in galls at 14 dpi**

**Table S4. Quantification of tomato transcripts in galls by RT-qPCR (G) relative to uninfected tomato roots (R), at 7 and 14 days post inoculation (dpi)**.

**Table S5. Gene ontology (GO) analysis of the genes DE in galls at 7 and/or 14 dpi**.

**Table S6. The number of raw reads from the three libraries obtained from galls and uninfected roots of *S. lycopersicum* at 7 dpi and 14 dpi**.

**Table S7. *De novo* prediction of *Solanum lycopersicum MIR* genes**.

**Table S8. MicroRNAs differentially expressed in galls at 7 and/or 14 dpi**.

**Table S9. 153 transcripts targeted by miRNAs in galls at 7 and/or 14 dpi identified by degradome sequencing and CleaveLand analysiss**.

**Table S10. Infection assays in *arf8 CRISPR* lines infected with *M. incognita* and comparison to WT**.

**Table S11. Genes differentially expressed between *crisprarf8* mutant galls and wild-type (WT) galls**.

